# Evolution of recombination suppression and sex determination on Y chromosomes of the plant genus *Mercurialis*

**DOI:** 10.64898/2026.03.31.715504

**Authors:** Jörn F. Gerchen, Daniel L. Jeffries, Stefan Grob, Victor Mac, John R. Pannell

## Abstract

Understanding why sex chromosomes repeatedly evolve recombination suppression, gene loss, and repeat accumulation remains a central challenge in evolutionary genomics. Plant sex chromosomes may be particularly informative, because they have often evolved recently from hermaphroditic ancestors. We studied the sex-linked region of the dioecious annual *Mercurialis annua* using new long-read genome assemblies of an XX female and a YY male, a published female assembly, linkage maps, and population-genomic data from several *Mercurialis* species. We identify two discrete nested evolutionary strata on the Y chromosome of diploid *M. annua*. A young stratum was generated by a large inversion and shows little degeneration, whereas an older stratum nested within it exhibits substantial gene loss, transposable-element accumulation, insertion of paralogous gene copies, and elevated X-Y sequence divergence. These findings indicate that recombination suppression evolved in at least two stages, with a recent inversion expanding an older non-recombining region. Comparative analyses among several *Mercurialis* species further show that the extent of sex-linked differentiation varies markedly among them. We also identify *APRR7* as the only gene showing consistent male-specific inheritance across the genus; this gene is a strong candidate master sex-determination gene. Together, our results refine the structure and gene content of the sex-linked region in *M. annua* and contribute to our understanding of the diversity of sex chromosomes in plants.

## Introduction

Sex chromosomes have evolved independently in various dioecious organisms, including animals, plants (Ming et al. 2011), algae (Coelho et al. 2019) and fungi (Branco et al. 2017). Many of these sex chromosomes share common properties, like the evolution of recombination suppression (Charlesworth 2017), the loss of functional genes (Skaletsky et al. 2003) and the accumulation of repetitive elements (Chalopin et al. 2015). In addition, sex chromosomes are generally thought to be important drivers of speciation, due to their disproportionate contribution to reproductive isolation (Qvarnström & Bailey 2009). However, the degree to which different sex chromosomes exhibit these common properties can vary considerably, and the underlying biological phenomena that cause these differences in sex-chromosome evolution are still poorly understood (Ponnikas et al. 2018). Specifically, it remains unclear what ultimately causes recombination suppression and gene loss, and whether similar outcomes of sex-chromosome evolution are caused by the same evolutionary processes in divergent species.

One property that varies considerately among species is the degree of recombination suppression and genetic degeneration of the sex chromosomes. Many sex chromosomes have large nonrecombining regions in which meiotic recombination is suppressed in the heterogametic sex. Recombination suppression is usually followed by genetic degeneration of the affected region, resulting in divergence between the homologous sequences that can range from microscopic differences to complete loss of one of the chromosomes. A common feature is the step-wise expansion of the length of the non-recombining regions, giving rise to so-called evolutionary strata with divergence between sex chromosomes and degeneration of the Y (or W) being a function of time since the origin of the respective stratum (Lahn & Page 1999). Nevertheless, although the degree of degeneration of a particular stratum and of the sex chromosomes in general are expected to increase with their age, several very old sex chromosomes in animals show little recombination suppression (Yazdi et al. 2014, Kuhl et al. 2021).

There are multiple non-mutually exclusive hypotheses for the expansion of recombination suppression on sex chromosomes. An adaptive explanation involves selection on so-called sexually antagonistic alleles, which increase fitness in one sex but reduce it in the other (Rice 1987). In such situations, we expect selection to favor suppressed recombination between a locus segregating for sexually antagonistic alleles and the sex-determining locus, strengthening the association between each respective allele and the sex in which it is beneficial. Alternative adaptive explanations involved the sheltering of recessive deleterious mutations due to fixed heterozygosity in the heterogametic sex—though recent analysis suggest that this scenario may apply only under limited circumstances (Charlesworth & Olito 2024). Neutral processes have also been explored (Jeffries et al. 2021), but it seems unlikely that they could apply except in very small populations.

It can also be difficult to disentangle cause and effect in the relationship between recombination loss and genetic degeneration on sex chromosomes. In some cases, the sex-determining locus lies in a chromosomal region that likely already had low recombination when it originated (Filatov et al. 2024, Li et al. 2026). This makes it unclear whether recombination suppression arose as a consequence of sex-chromosome evolution or predated it (Rifkin et al. 2020). Furthermore, nonrecombining regions on sex chromosomes (Śliwińska et al. 2016), but also other parts of the genome such as pericentromeric regions, often show a strong accumulation of transposable elements (Bousius et al. 2025). The spread of transposable elements is thought to be shaped by multiple factors, including recombination, which can remove TE copies (Kent et al. 2017), and the methylation-mediated silencing of active elements (Zhou et al. 2020). TEs may therefore contribute to sequence divergence and increase the rate of structural variation through mechanisms other than recombination alone. As such, they could promote the expansion of recombination suppression (Hobza et al. 2015; Huang et al. 2025), thereby favouring further TE accumulation in a positive feedback loop.

Plant sex chromosomes provide particularly valuable systems for studying the early stages of sex-chromosome evolution and, potentially, for disentangling cause and effect in the evolution of recombination suppression. In dioecious plants, separate sexes and sex chromosomes have frequently evolved de novo from hermaphroditic ancestors (Westergaard 1958; Charlesworth 2016). The canonical model for the evolution of separate sexes in plants posits the evolution of recombination suppression between two loci: one carrying a male-sterility allele, resulting in females, and the other carrying an allele that promotes male fertility while reducing female fertility, resulting in males (Charlesworth & Charlesworth 1978). In this framework, recombination suppression is viewed as part of the initial process by which separate sexes and sex chromosomes evolve. While several plant species provide strong evidence for two linked loci with complementary sterility effects in the two sexes (Akagi et al. 2016; Harkess et al. 2020; Massonnet et al. 2020), other systems show patterns of sex determination that may have arisen through different evolutionary routes, such as successive regulatory changes in sex expression or epistatic interactions with autosomal genes (Akagi et al. 2014), although even in these cases an initial two-mutation pathway to dioecy cannot be excluded.

The plant genus *Mercurialis* (Euphorbiaceae) provides a particularly interesting system for studying the evolution of plant sex chromosomes, especially against a background of recent transitions between combined and separate sexes and variable ploidy (Pannell et al. 2004). Although dioecy is ancestral in *Mercurialis*, several species in a clade of annual taxa have undergone transitions either to monoecy or to the rare sexual system androdioecy, in which males coexist with hermaphrodites (Pannell 1997). Previous analyses indicate that males in both the annual clade and its perennial sister clade share a common ancestral XY system of sex determination (Gerchen et al. 2022). Interestingly, the Y chromosome in hexaploid *M. annua* appears to have introgressed from a more distantly related perennial species (Gerchen et al. 2022). Two previous studies estimated the size of the non-recombining region of the Y chromosome in dioecious, diploid *M. annua* to be between 14.5 and 19 Mb, containing approximately 500 genes (Veltsos et al. 2018; Veltsos et al. 2019). One study used bacterial artificial chromosome (BAC) sequences anchored to the male-specific linkage map, whereas the other inferred gene content from transcripts mapping to the male-specific region of the linkage map. Both approaches have limitations because they rely on linkage-map information rather than direct physical genome assemblies. As a result, they may overestimate the size and gene content of the non-recombining region. These inferences therefore require validation using chromosome-level genome assemblies together with population-genomic data.

Here, we present an improved inference of the size and gene content of the non-recombining region in *M. annua* based on new long-read genome assemblies of both an XX female and a male carrying two Y chromosomes. We complement these assemblies with a range of published and unpublished population-genomic datasets from different *Mercurialis* species, as well as an additional linkage map for male *M. annua*. We report evidence of two clearly distinguishable evolutionary strata on the Y chromosome of *M. annua*: one young stratum that has been generated by a large chromosomal inversion and that shows otherwise very little evidence of degeneration and sequence divergence; and an older stratum nested inside the younger one, which shows substantial evidence of gene loss, accumulation of transposable elements and insertion of paralogous gene copies. Population genomic evidence from further *Mercurialis* species reveals that the extent of recombination suppression can vary significantly between species. We also propose a candidate master sex determination gene called APRR7 located in the old stratum.

## Material and Methods

### Genome assemblies

We used three genome assemblies. Two assemblies were newly generated assemblies of a male (the ‘YY assembly’) and a female (the ‘XX assembly’), both of which were offspring of a single self-fertilized XY male whichthat was partially feminized by external application of cytokinin (Li et al. 2019). We used a restriction enzyme-based assay to identify YY males (Li et al. 2019) and used a single YY male and an XX female issuing from the same cross. The other assembly was a previously published assembly of an unrelated female from the Darwin Tree of Life Project (Christenhusz et al. 2024), (the ‘XXdtol assembly’). We used all three assemblies for comparative analysis of gene content in the sex-linked region, allowing us to identify potential artifacts introduced by our assembly and annotation pipeline; note that the XX and YY assemblies were generated and annotated using the same protocol, which involved Oxford Nanopore longread sequencing and annotation using Braker 3 (Gabriel et al. 2024), whereas the XXdtol assembly was based on PacBio Hifi sequencing and was annotated using the Ensembl genebuild pipeline for non-vertebrate species (https://rapid.ensembl.org/info/genome/genebuild/anno.html).

### Whole-genome sequencing

For the new assemblies of the XX female and the YY male, we undertook two rounds of high molecular weight DNA extraction from each individual using two different protocols. First, we extracted high molecular weight DNA from extracted nuclei. For this, we first ground 5 grams of fresh leaves in liquid nitrogen and added 50 mL isolation buffer (15 mM Tris, 10 mM EDTA, 130 mM KCI, 20 mM NaCI, 8% (m/V) PVP-10, 0.1% Triton X-100, 7.5% (v/v) 2-Mercaptoethanol, pH 9.4). We filtered the nuclei suspension through 50 μL mira-cloth, and we then added 500 μL Triton X-100, which we pelleted by centrifuging at 2000g at 4 °C for 10 min. We further added 10 mL of Carlson lysis buffer (100 mM Tris pH 9.5, 2% CTAB, 1.4 M NaCl, 1% PEG 6000, 20 mM EDTA, prewarmed to 74°C) and 25 μL of 2-Mercaptoethanol and incubated the solution at 74°C for 2 h.

We extracted samples twice by adding 1.0 volume of 24:1 chloroform/isoamyl alcohol to the aqueous phase and centrifuging at 5,000 g for 10 min at 4°C. We added 0.1 volume of 3 M NaOAc and precipitated DNA by adding 1.0 volume of isopropanol and incubating overnight. After centrifuging at 4,500 g for 90 min, we washed pellets by adding 10 mL of 70% EtOH and centrifuging at 4,500 g for 90 min. We discarded the supernatant and air-dried pellets for 10 min and resuspended them in Qiagen G2 buffer. We added 2 μL/mL RNase A (Qiagen, 100 mg/mL) to the resuspended DNA pellets, mixed and incubated for 5 min at room temperature and added 10 μL/mL Proteinase K. We incubated the sample at 50°C for 1 h and cleaned it using Qiagen Genomic Tip 500/G columns, following standard instructions. We added 10.5 mL of isopropanol at room temperature to the eluted DNA and precipitated the DNA overnight. We then centrifuged at 4,500 g for 90 min at 4°C, discarded the supernatant and added 40 mL of ice-cold 70% EtOH, centrifuged at 4,500 g for 30 min at 4°C, discarded the supernatant, let the pellet air dry for 10 min and resuspended the DNA in 10 mM TrisCl (pH 8).

For a second high molecular weight DNA extraction, we ground 5 g of fresh leaves in liquid nitrogen and added 40 mL of Carlson lysis buffer with 2.5% (v/v) 2-Mercaptoethanol and 80 μL of RNase A (Qiagen, 100 mg/mL). We incubated samples at 65°C for 1 h and, after cooling to room temperature, we extracted DNA by adding 1.0 volume of 24:1 chloroform/isoamyl alcohol to the aqueous phase and centrifuging at 3,500 g for 15 min at 4°C, retaining the aqueous phase. We precipitated DNA by adding 0.7 volumes of isopropanol and incubating at -80°C for 15 min. We centrifuged the sample at 3,500 g for 15 min at 4°C and discarded the supernatant. We resuspended the pellet in Qiagen G2 buffer and cleaned it using Qiagen Genomic Tip 500/G columns following standard instructions. We added 0.7 volumes of isopropanol to the eluted DNA and precipitated DNA at -20 °C overnight and centrifuged at 3,500 g for 45 min at 4°C. We discarded the supernatant and washed the pellet by adding 10 mL of ice cold 70% EtOH and centrifuged at 3,500 g for 10 min at 4°C. We discarded the supernatant let the pellet air dry for 10 min before resuspending it in 10 mM TrisCl (pH 8).

We sequenced genomic DNA on Oxford Nanopore flow cells (v. 9.4.1) using an Oxford Nanopore Minion. Libraries were prepared using either ligation sequencing kit (SQK-LSK109) following standard instructions or the rapid sequencing kit (SQK-RAD004) from Oxford Nanopore, following standard instructions. Finally, we also sent high molecular weight DNA from both plants to Novogene UK, who generated additional sequencing libraries using Oxford Nanopore Promethion. In addition, we sent genomic DNA from both samples for short read sequencing on the Illumina NovaSeq S6 platform with 150 bp paired-end reads to Novogene UK.

### Generation of a HiC scaffolding library

We used 5 grams of fresh leaves of the individual used for the YY assembly. We used the HiC protocol described in Grob et al. (2022) with the HindIII restriction enzyme and generated Illumina libraries using Roche KAPA Hyperprep library kits and had them sequenced using Novaseq S4 PE150 by Novogene UK.

### Genome assembly and annotation

Raw Oxford Nanopore reads from self-generated libraries were base-called using Guppy 4.2.2 (Oxford Nanopore). ONT reads from all described methods were used to generate genome assemblies using Canu 2.2 (Koren et al. 2017). We ran one round of polishing on the raw genome assemblies using the Oxford Nanopore long reads with Racon (Vaser et al. 2017). For this, we aligned ON reads against the genome assemblies using Minimap2 (Li, 2018) and ran Racon 1.4.3 (Vaser et al. 2017) with standard parameters. We then ran a second round of polishing using both short and long-reads using HyPo 1.0.3 (Kundu et al. 2019). For this, we aligned both the long reads and the short Illumina reads using minimap2. We removed haplotypic duplications using the purge_dups 1.2.6 pipeline (Guan et al. 2020). For this, we aligned ON long reads against the polished assemblies using Minimap2 (Li, 2018) and ran the purge_dups pipeline with default parameters. We scaffolded both genome assemblies using yahs (Zhou et al. 2023) by aligning HiC libraries using the arima pipeline (https://github.com/ArimaGenomics/mapping_pipeline). For the YY male, we used our self-generated HiC data, while for our assembly of the XX female we used published HiC data from the Darwin tree of life project. We then manually assessed our scaffolded assemblies using JuiceBox and corrected the orientation of inverted sequences as well as separated merged chromosome scale scaffolds. We generated a de-novo repeat library from the YY assembly using EarlGrey 5.1.0 (Baril et al. 2024) using five iterations. We used the YY repeat annotation to annotate repeat elements on our XX assembly as well as the female genome assembly published by the dtol project using EarlGrey.

We removed contaminations from our genome assemblies by using the taxonomy module in mmseqs2 16.747c6 (Steinegger et al. 2017) to find homology between contigs and the Uniprot database, and we assigned taxonomic assignments to individual sequences. We then used custom Python scripts that employed the taxonomy functions of the ete3 3.1.3 package (Huerta Cepas et al. 2016) to infer taxonomic class from the mmseqs results where possible, together with an additional script to remove DNA sequences from the assemblies that were assigned to classes other than angiosperms. We then assessed the completeness of our assemblies using BUSCO 6.0.0 (Tegenfeld et al. 2025) using the malpighiales_odb12 lineage dataset.

We generated new gene annotations for our new YY and XX assemblies with Braker 3.0.8 (Gabriel et al. 2024), using softmasked genome assemblies and both rnaseq data from multiple tissues and life stages and protein data from the Viridiplantae subset of Orthodb v. 12 (Tegenfeld et al. 2024).

### Analysis of synteny between genome assemblies

We assessed orthology of gene models across annotations and synteny between assemblies using GENEspace 1.3.1 (Lovell et al. 2022), based on the results of Orthofinder (Emms et al 2018). To generate input for GENEspace, we used the agat_sp_keep_longest_isoform.pl script from AGAT 0.8.1 (Dainat 2022) to identify the longest isoforms for all three annotations, and we then used the agat_sp_extract_sequences.pl script with the –p option to extract amino acid sequences and used the agat_convert_sp_gff2bed.pl with the —sub option set to “gene” to convert gff files to bed files. We used grep –v “transcript” to remove duplicate entries for transcripts. Based on riparian plots generated by GENEspace, we identified large structural variations and focused on the largest inversion on the YY assembly, which we identified as the male-specific non-recombining region.

### Manual curation of gene models in the sex-linked region

A key step was to improve the reliability of the gene annotations generated for the XX and YY assemblies as well as the automated annotation by the automated NCBI annotation pipeline in the sex-linked region, as well as to annotate gene models. For functional annotation and inference of orthology between the three genome assemblies and other angiosperm species, we ran Orthofinder 3.1.1 (Emms et al. 2018) on the gene models from the three genome assemblies as well as protein models based on twelve genome assemblies from the NCBI RefSeq project (Table S1). Based on this analysis, we further used the protein alignments generated by Orthofinder to manually check the orthology of alignments and the gene trees to test if gene models form the three *M. annua* genome assemblies form a monophyletic clade. For annotation, we used the annotation from the *M. annua* XXdtol assembly also for the other two genome assemblies if they formed a monophyletic clade, except for cases when the *M. annua* genome assembly was annotated as unknown protein;! in such cases, we sought the most common annotation of the other genome assemblies from the same orthogroup.

For further manual curation of gene models, we used Apollo 2.8.1 (Dunn et al. 2019), and we generated additional ab-initio gene predictions for all three genome assemblies using helixer 0.3.4 (Holst et al. 2025) and tiberius 1.1.7 (Gabriel et al. 2024b) to provide alternative gene models. Where genes were missing from one of the annotations generated by BRAKER, we used these alternative annotations. In addition, we aligned RNAseq data of male samples against the YY genome assembly using STAR 2.7.10b (Dobin et al. 2013) and we used Megadepth 1.2.0 (Wilks et al. 2021) to estimate sequencing depth from alignments and output it in wig format, which we also used in Apollo as additional evidence of gene expression. We further used stringtie 3.0.3 (Pertea et al. 2015) to assemble transcript models and added them to Apollo. We then manually inspected all gene models from the SDR and sought to construct consensus sequences between all three genome assemblies, with the aim to cover as much orthologous sequence as possible. At the same time, we tried to identify and exclude transposable elements, which were erroneously annotated as genes. We typically did this to gene models which had a high copy number in the orthogroup, were short and had few exons, overlapped with regions marked as repeats by EarlGrey and which were annotated as unknown proteins in all annotations within the orthogroup.

### Evolutionary rates and analysis of deleterious variants

We conducted pairwise alignments of the amino acid sequences from the SDR based on the curated genome annotations for the three genome assemblies using MAFFT 7.525 (Katoh et al. 2013) with the --localpair option enabled and the --maxiterate option set to 1000. We then used translatorX 1.1 (Abascal et al. 2010) to generate codon-based CDS alignments based on the amino acid alignment. Finally, we used the yn00 model (Yang & Nielson 2000) of PAML 4.10.10 (Yang 2007) to estimate pairwise synonymous and non-synonymous substitution rates (dS and dN, respectively).

We used a complementary approach to quantify the effects of variation in coding sequences in the SDR. We assessed the effects of variants from the YY and XX sequencing data on our hand-curated gene models from the XXdtol assembly in the SDR. We aligned short read data from both the XX and YY individuals against the Xxdtol genome assembly using BWA mem 0.7.18 (Li 2013) and called variants using freebayes 1.3.8 (Garrison & Marth 2012). Since we were only interested in the SDR and since both the XX and the YY individuals were the offspring of a single selfed XY male, we assumed that both the XX and the YY individuals inherited the twice the same, unrecombined SDR region and we set the ploidy for both individuals to haploid. We then filtered the resulting VCF file to include only variants with a QUAL score > 100 and we used variant effect predictor (VEP) 115.2 (McLaren et al. 2016) to classify variants based on their effects on coding sequences.

### Population genomic analysis of diploid *M. annua*

For population genomic analyses, we used a previously generated exon-capture dataset of 20 male and female individuals of *M. annua* from across the species range in central and southeastern Europe (González-Martínez et al. 2017). Since this dataset was designed by including all non-repetitive sequences of an early draft assembly of an XY male *M. annua* genome, it included probes at high density, including those located on both the Y and X chromosomes.

We used a custom snakemake (Köster & Rahmann 2012) workflow implemented in the freebayes_single git branch, to align reads of both sexes against the YY and XX_dtol genomes and call variants. We trimmed single-end reads using trimmomatic 0.39 (Bolger et al. 2014) with default parameters. We then aligned trimmed reads against the reference genome using BWA mem 0.7.18 (Li 2013) and marked PCR and optical duplicates using Picard MarkDuplicates 3.3.0 (Broad Institute 2019). We inferred sequencing depth at the sites of the original exon-capture probes, by extracting the sequences of the original exon-capture probes from the draft genome assembly of *M. annua* using bedtools 2.31.1 (Quilan 2014) and aligning their sequences against the YY and XX_dtol genomes using Minimap2 (Li 2018). We extracted probes with an alignment quality of 60 from the resulting paf file and inferred the sequencing depth of bam files at the locations using PanDepth 2.25 (Yu et al. 2024). We then used freebayes 1.3.8 (Garrison & Marth 2012) to call variants at both variant and invariant sites (using the --report-monomorphic option), followed by filtering using bcftools 1.21 (Danecek et al. 2021). We first used the filter module of bcftools to set all genotypes with a smaller than 4 to no-call and used a custom python script to remove sites, which had more than 50% missing data in either males or females. We then used the fill-AN-AC plugin of bcftools to recalculate allele counts and used the -a option of bcftools view to remove genotypes which were not present anymore after the previous filtering steps. We then subset biallelic snps using the -m2, -M2 and -v snps parameters of bcftools view and removed variants with a QUAL score smaller than 100. We then subset invariant sites with the -M1 parameter of bcftools view and merged quality filtered biallelic SNPs and invariant sites using bcftools concat.

### Linkage mapping

We used a modified version of our snakemake workflow (implemented in the freebayes_rnaseq git branch) to align RNAseq data published in Veltsos et al. (2019) against the XXdtol genome assembly and to call variants. We used STAR (Dobin et al. 2013) with the options SortedByCoordinate activated and twopassMode set to basic to align RNAseq reads. We then used the SplitNCigarReads module from GATK to split alignments. We used GATK HaplotypeCaller 4.6.10 (Poplin et al. 2017) to call variants for individual samples using the HaplotypeCaller module with the —dont-use-soft-clipped-bases option enabled. We then we used the GenomicsDB module to build a common Genomics DB for all samples and used the GenotypeGenomicsDB module for combined genotyping. We filtered variants to include only sites found in the annotated gene models for the XXdtol genome assembly, and we removed variants that were found in only a single individual.

We used LepMap3 0.5 (Rastas 2017) to build male and female linkage maps based on the genotyped RNAseq data. We used the ParentCall2 function with the options removeNonInformative and halfSibs activated to build a parent call file, from which we then used the OrderMarkers2 function with the minError parameter set to 0.01 to account for genotyping errors and set improveOrder to 0 to set the gene order based on the reference genome assembly to build separate linkage maps for each chromosome.

### Male linkage map

We built a new combined male linkage map for *M. annua* from three interspecific F_1_ crosses between a male *M. annua* and three female *M. huetii*.

We extracted DNA using either a robotic BioSprint 96 workstation (Qiagen) and the BioSprint DNA Plant kit (Qiagen) or using the DNeasy Plant Mini kit (Qiagen). We quantified extracted DNA using the Qubit dsDNA HS Assay Kit on a Qubit fluorometer.

We generated Genotype data using a modified ddRad protocol (Peterson et al. 2012). For each sample, we first digested about 100 ng DNA diluted to 22 μL by adding 2.5 μL Smartcut buffer (NEB), 0.4 μL EcoRI-HF (NEB) and 0.4 μL Taq1 (NEB) restriction enzymes and incubating at 37°C for 30 min, 65°C for 30 min and 80°C for 20 min.

We then ligated P1 and P2 adapters described in Peterson et al. 2012, which were diluted to 40 μM in 1x annealing buffer (50mM NaCl, 10 mM). We added 3 μL rATP (10 mM), 2 μL annealed P2 adapter (3 μM), 0.8 μL 10x T4 ligase buffer (NEB) 1 μL T4 ligase (400 U/μL, NEB), 22 μL digested DNA and 2 μL annealed P1 adapter (0.3 μM) and incubated for 20 min at 23°C followed by 10 min at 65°C. We then size selected 300 μL of pooled samples for each library to 550 bp. For this we first added 0.57x volume of Ampure XP beads incubated for 10 min at room temperature and saved the supernantant, to which we added 0.12x volume of Ampure XP beads (Beckman Coulter), incubated on magnetic stand, washed the beads with 70% ethanol and eluted DNA in 30 μL water for 2 min.

We selected for biotin-labeled P2 adapters using M-270 Dynabeads (Invitrogen). We washed 15 μL Dynabeads 3 times in 1x bind and wash buffer (5 mM Tris-HCl pH 7.5, 0.5 mM EDTA, 1M NaCl), resuspended them in 30 μL 2x bind and wash buffer, added the size selected DNA and incubated for 15 min at room temperature. We discarded the supernatant, washed the beads three times using 1x bind and wash buffer and resuspended the beads in 45 μL water.

We then amplified size selected fragments by PCR. We used single indexed primers recommended by the modified ddRAD protocol (Peterson et al. 2012). We added 45 μL bead suspension, 3 μL forward primer (10μM), 3 μL reverse primer (10 μM) and 50 μL KAPA HiFi Hotstart Ready Mix (Roche). PCR program was 2 min at 95 °C initial denaturation and 11 cycles at 98 °C for 20 s, 65 °C for 20 s and 72 °C for 30 s.

We cleaned PCR products in 0.7x Ampure XP beads according to manufactorers instructions and eluted PCR products in 20 μL water. Libraries were sequenced using 150 bp single end reads by the Genomics Technologies Facility (GTF) at the University of Lausanne.

We demultiplexed and quality filtered raw Illumina reads using Stacks 2.68 (Catchen et al. 2013). For this study, we were interested in the male *M. annua* linkage map; we thus reconstructed alleles that were specific to the *M. annua* fathers. To this end, we ran a modified version of the Stacks pipeline. We first ran the ustacks program on all samples, allowing a maximum of four nucleotides between stacks. Then we ran cstacks and build the allele catalogue using only the fathers from all families, allowing a maximum of four mismatches between samples. We then ran sstacks module to match reads from all samples against the catalog and we ran tsv2bam to transpose read data. We then extracted the names of reads from the resulting bam files using samtools 1.18 (Danecek et al. 2021) and used a custom python script based on biopython 1.70 (Cock et al. 2009) to subset our original reads to include only those that matched to paternal genotypes. We then aligned the subset of reads against the YY assembly using bwa mem 0.7.18 (Li 2013) and used the gstacks module with a minimum mapping quality of 40 to call snps and the populations module to filter sites, which were found in at least 60% of individuals in at least one population and with haplotype wise filters applied. We then used a custom python script to generate modified VCF files for linkage mapping. This script filtered snps with a minor allele frequency smaller than 0.2, replaced missing or homozygous paternal genotypes with heterozygotes based on the alleles found in their offspring, set heterozygous offspring genotypes to nocall and added a dummy reference allele to the remaining homozygous offspring genotypes. We then used the modified VCF file to build a linkage map using LepMap3 0.5 (Rastas 2017). We first used the ParentCall module with the removeNonInformative and halfSibs parameters activated. Then we used the OrderMarkers2 module for each chromosome, using the order of sites from the VCF file and the options improveOrder deactivated and proximityScale set to 200.

### Population genomic analyses of other *Mercurialis* species

We used a previously generated exon-capture dataset (Gerchen et al. 2022), which included between 5 and 10 samples for each sex (either males and females, or males and monoecious individuals) for other annual and perennial species with different ploidies. We used a different branch from our custom Snakemake pipeline to align the paired-end reads against the YY genome assembly and to call variants and infer sequencing depth at probes, which was performed similarly to the single-end version described above, with the exception that, here, bwa mem was run for paired-end reads.

For diploid *M. annua* and *M. huetii* we estimated *F*_ST_ between male and female samples in 50 kb windows using pixy 2.0.0.beta8 (Korunes & Samuk 2021). For the other allopolyploid species, we were unable to use *F*_ST_ due to the presence of fixed heterozygotes in allopolyploids. Instead, we inferred *F*‘_ST_, which was specifically developed for estimating differentiation in allopolyploids and which is based on allelic phenotypes (Obbard et al. 2006). We developed a custom python script, which infers *F*‘_ST_ in sliding windows, and we inferred *F*‘_ST_ between either males and females, or males and monoecious individuals (for hexaploid *M. annua*), in 50 kb windows, including variants for which fewer than 50% of samples of each sex had missing data.

We generated pooled sequencing libraries for male and female samples of *M. elliptica* and for male and monoecious samples of hexaploid *M. annua*. We extracted DNA from dried leaves using the DNEasy blood and tissue kit for hexaploid *M. annua* and a robotic BioSprint 96 workstation (Qiagen) and the BioSprint DNA Plant kit (Qiagen) for *M. elliptica.* We estimated DNA concentrations using a Qubit DNA kit and pooled samples in equimolar concentrations. We then had sequencing libraries prepared and samples sequenced using Illumina 150 bp paired-end reads using a PCR-free protocol by Novogene Ltd. We aligned reads against the YY genome assembly using bwa mem 0.7.18 (Li 2013) and generated a new GFF file, which contained our hand-curated gene models for the region inside the SDR and the uncurated longest gene models from the BRAKER annotation outside the SDR. We used PanDepth 2.25 (Yu et al. 2024) to estimate the sequencing depth at the CDS of each gene model and calculated an adjusted mean depth per gene model for each pool by multiplying the depth times with the proportion of the gene covered.

## Results

### Genome assemblies

The two new *M. annua* genome assemblies had total lengths of 420,999,989 bp and 452,796,065 bp, assembled into 539 and 507 scaffolds, with N50 values of 43,150,279 bp and 39,860,971 bp for the YY and XX assemblies respectively. Compared with the previously generated XXdtol assembly, which has an assembly size of 453,773,921 in 63 scaffolds and an N50 value of 56,051,264 Mb, the new assemblies here are moderately less contiguous, which mostly results in smaller assembled centromeric regions (Fig 1 D). Initial runs of BRAKER identified 21,665 and 22,475 gene models on the YY and XX assemblies, respectively. BUSCO completeness values were 99.3% (98.2% single copy, 1.1% duplicated, 0% fragmented and 0.7% missing) and 99.8% (98.9% single copy, 0.9% duplicated, 0% fragmented and 0.2% missing) for the YY and XX assemblies respectively, compared to 99.9% (99.5% single copy, 0.5% duplicated, 0% fragmented and 0.1% missing) for the XXdtol assembly. These values imply a very high level of completeness and low levels of duplication in both the new genome assemblies. Comparison of gene models using GENESPACE shows that the gene content in the chromosomal scaffolds of our assembly is largely consistent and syntenic with the XXdtol assembly (Fig. S1).

**Figure 1.**
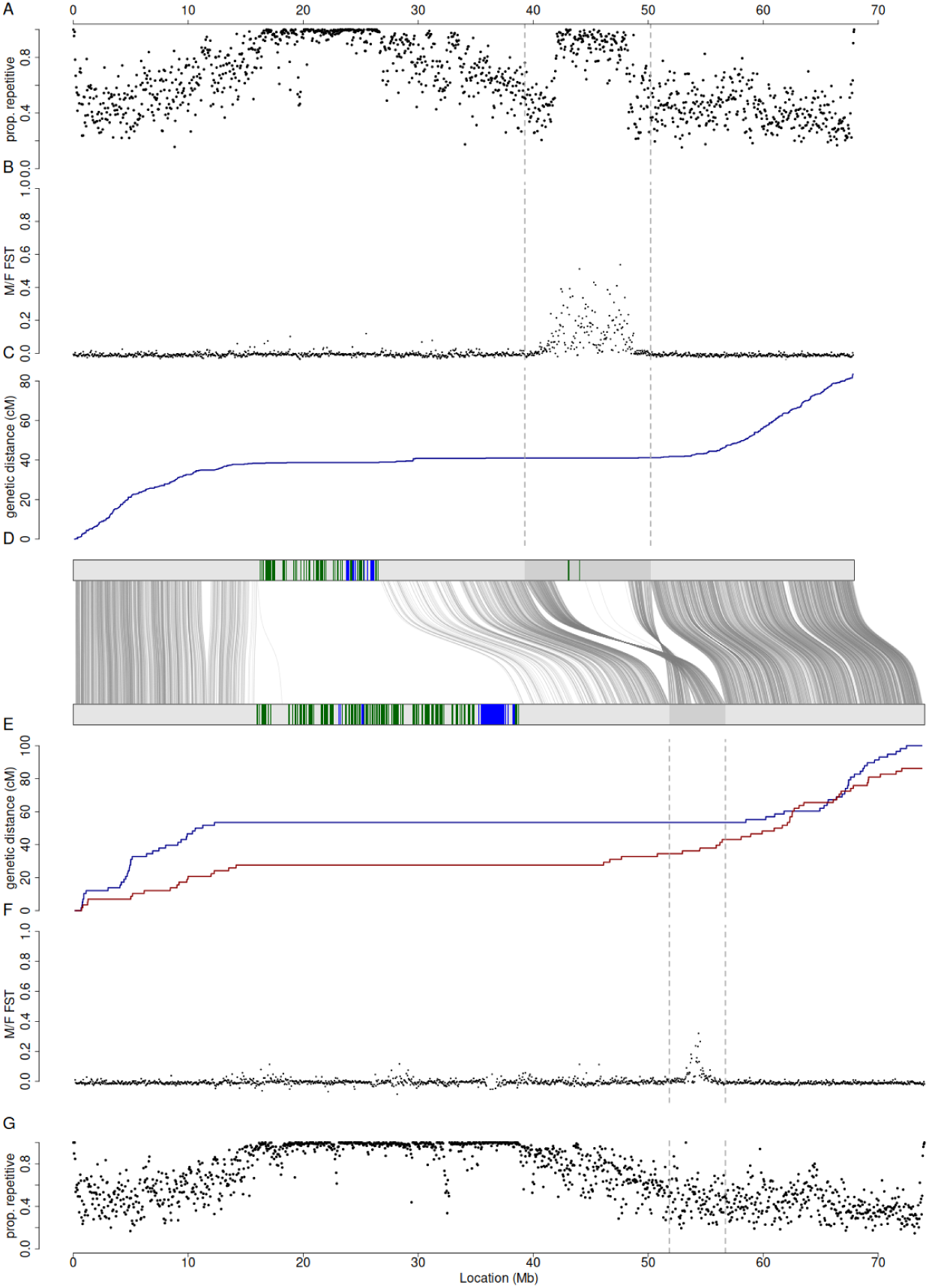
Plots for Chromosome 1 on the YY and XXdtol genome assemblies. A: proportion of repetitive sequences identified on the YY assembly. B: mMale-fFemale *F*_ST_ based on exon-capture dataset. C: male linkage map based on cross between an *M. annua* male and *M. huetii* females. D: synteny between shared gene models on the YY (top) and XXdtol genome assemblies (bottom). Green and blue areas indicate regions enriched for the two most common centromeric tandem repeats (green: 60bp repeat, blue: 99bp repeat), the darker grey regions highlights the large inversion on ChrChromosome 1, curved grey lines show locations of syntenic genes found in all three genome assemblies. E: female (dark red) and male (blue) linkage maps based on RNAseq data from Veltsos et al. (2019). B: male-female *F*_ST_ based on exon-capture dataset. G: proportion of repetitive sequences identified on the XXdtol assembly.

Based on the GENESPACE analysis, we identified four large inversions between the three genome assemblies used in our analysis. Two inversions were found on Chromosome 1 of the YY assembly, and the other two were found on Chromosome 4 and Chromosome 7 of the XXdtol assembly (Fig. S1). One of the inversions on Chromosome 1 was the largest, spanning 10,943,025 Mb on the YY assembly, while the equivalent regions of the XXdtol and XX assemblies were only 4,877,125 bp and 4,826,375 bp, respectively, implying that the inverted region on the Y chromosome is approximately 6 Mb larger than on the X chromosome. We annotated 69.4% of the YY assembly, 71.5% of the XX assembly and 71% of the XXdtol assembly as repetitive. Besides putative pericentromeric regions (see below), we identified one region with strongly increased repeat density inside the largest inversion on Chromosome 1 on the YY assembly but not on the XX or XXdtol assemblies (Fig. 1A, Fig 1G). We identified two common putative centromeric repeats with lengths of 60 and 99 bp using TRASH, and we identified large regions enriched for these repeats on all eight chromosomes in all three genome assemblies, which also corresponds to regions of increased repeats identified by Earl grey. These regions were located approximately in the centre of seven out of the eight chromosomes, with the exception of Chromosome 4, where it was located on the edge.

### Linkage maps

We generated three linkage maps based on alignments of genotyped offspring from crosses to either the YY or XXdtol genome assemblies. We generated a male and a female linkage map by aligning the RNAseq data generated by Veltsos et al. (2022) against the XXdtol assembly, which had a total length of 629.31 cM and 748.28 cM for the female and male maps, respectively. We also generated a male linkage map by aligning RADseq data of a cross between a male *M. annua* and a female *M. huetii* against the YY assembly, which had a total length of 433.66 cM. In all crosses, we identified large regions of reduced recombination at putative pericentromeric regions. In addition, we identified strong differences between male and female recombination maps on Chromosome 1 (Fig. 1C, Fig. 1E). On the cross based on the smaller set of RNAseq data, the non-recombining pericentromeric region in males extends much further than in females, so that it encompasses also the large inversion found on the YY assembly (Fig. 1E). In contrast, in females there is still substantial recombination within and before the inverted region. In the male linkage map based on the larger cross, there was also no evidence for recombination found within the inversion. However, unlike the RNAseq map, this linkage map suggests a small number of recombinants between the centromere and the inverted region (Fig. 1C).

### Population genomics of diploid *M. annua*

Our population genomic analysis of exon-capture data from 20 males and 20 females of diploid *M. annua* revealed a strong peak of high male-female *F*_ST_ in the centre of the large inversion of Chromosome 1, which encompassed a much larger region on the YY assembly than on the XXdtol assembly (Fig. 1B, Fig. 1F).

### Gene content of the SDR region

After manual curation, we retained a total of 412 gene models in the non-recombining regions of both the XX and XXdtol genome assemblies, and 401 gene models in the non-recombining region of the YY genome assembly. Based on the presence and absence of orthologous genes, the peak in male-female *F*_ST_, and the accumulation of transposable elements, we could split the nonrecombining region into two discreet regions, which we label the younger and the older stratum (Fig. 2A). The younger stratum contains a total of 387 gene models, which are found on all three genome assemblies. The older stratum has divergent gene content between the YY assembly and the XX and XXdtol assemblies. Twelve genes were found on the YY assembly, but were missing from both the XX and XXdtol assemblies. Based on gene trees generated by Orthofinder, we could determine the putative origin of nine of them, with seven being paralogous copies of other genes found in the new stratum and two being paralogous copies of autosomal genes from Chromosomes 2 and 8. Twenty-two genes were found on both the XX and XXdtol assemblies, but missing from the YY assembly, and three genes from in the old stratum were shared between all three genome assemblies.

**Figure 2:**
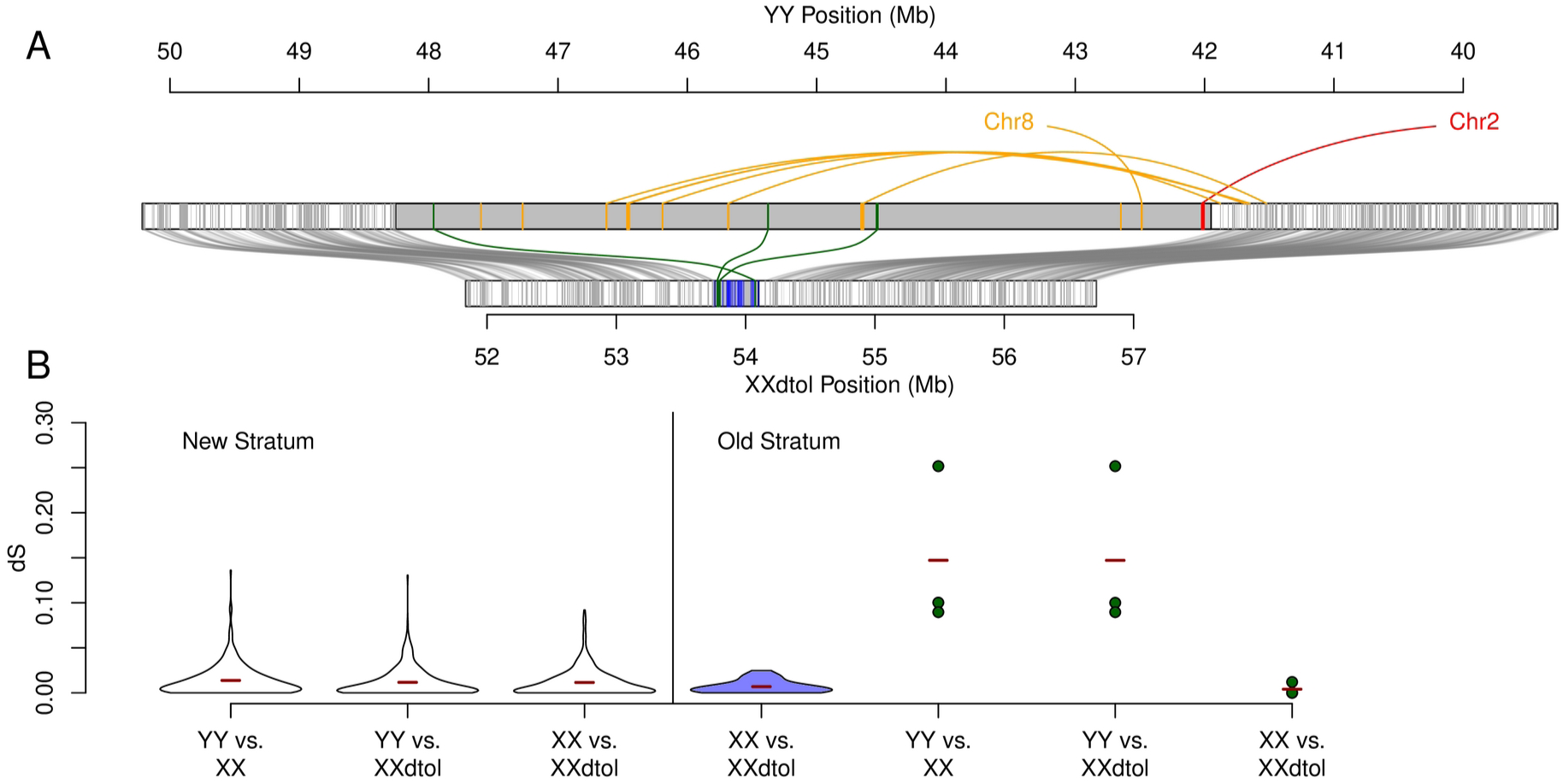
Gene content of the SDR and synomymous divergence between sex-linked genes. A: Comparison of the sex-linked regions on the YY (top) and XXdtol (bottom) assemblies. Note that the inversion on the YY assembly has been de-inverted. Grey blocks and curved lines indicate syntenic genes between X and Y chromosomes in the new stratum. The old stratum is indicated in dark grey color. Orange and red genes indicate genes specific to the YY assembly, the position marked in red indicates the position and origin of the putative master sex determination gene. Curves on the top of the YY assembly indicate the putative origin of YY specific genes from either the new stratum or other chromosomes. Green blocks/curves indicate genes on the old stratum shared between X and Y chromosomes, and blue blocks indicate the location of genes specific to the X chromosome. B: Synonymous substitution rate (dS) for pairwise comparisons of genes from all three assemblies found on the new stratum (left part) and the old stratum (right part). Violin plots indicate the distribution of dS values, red bars indicate the mean. Note that since only three genes were shared between the X and Y chromosomes no violin plots, but individual values were plotted.

We identified 16 premature stop codons in genes found on both the X and Y chromosome based on the YY reads aligned against the XXdtol assembly, but only one premature stop codon based on XX reads. In addition, we identified 22 frameshift variants based on YY reads and twelve frameshift variants based on XX reads. In total, frameshift variants affected 27 XXdtol gene models based on YY reads and eleven gene models based on XX reads (Suppl. Table 1).

In pairwise comparisons, the mean synonymous substitution rates, dS, in the new stratum were 0.0138, 0.0116 and 0.0115 for comparisons between the YY and XX assemblies, between the YY and XXdtol assemblies and between the XX and XXdtol assemblies, respectively. In the old stratum, mean dS was 0.00686 for comparison of X-hemizgous genes between the XX and XXdtol assemblies. The mean dS for the three genes retained as both Y and X copies on the old stratum were 0.147, 0.147 and 0.004 for comparisons between the YY and XX assemblies, between the YY and XXdtol assemblies and between the XX and XXdtol assemblies, respectively (Fig. 2B).

### Sex-linked variantion in other *Mercurialis* lineages

Our results showed that the degree of sex-chromosome differentiation between sexes differs between *Mercurialis* species. While diploid *M. annua* shows a strong signal of male-female *F*_ST_ in the old stratum (Fig. 3A), there is a much larger region of increased male-female *F*_ST_ in *M. huetii*, which covers a region that begins approximately 15 Mb further to the right of the SDR in diploid *M. annua* (Fig. 3B). In *M. canariensis*, peaks of male-female *F’*_ST_ are also found in the new stratum, but are otherwise limited to the SDR region of *M. annua* (Fig. 3C). In hexaploid *M. annua* and the three perennial *Mercurialis* species, no striking peaks of differentiation are observed. *F’*_ST_ between males and females, or between males and monoecious individuals in the case of hexaploid *M. annua,* is only moderately increased in the SDR (Fig. 3D-G).

**Figure 3.**
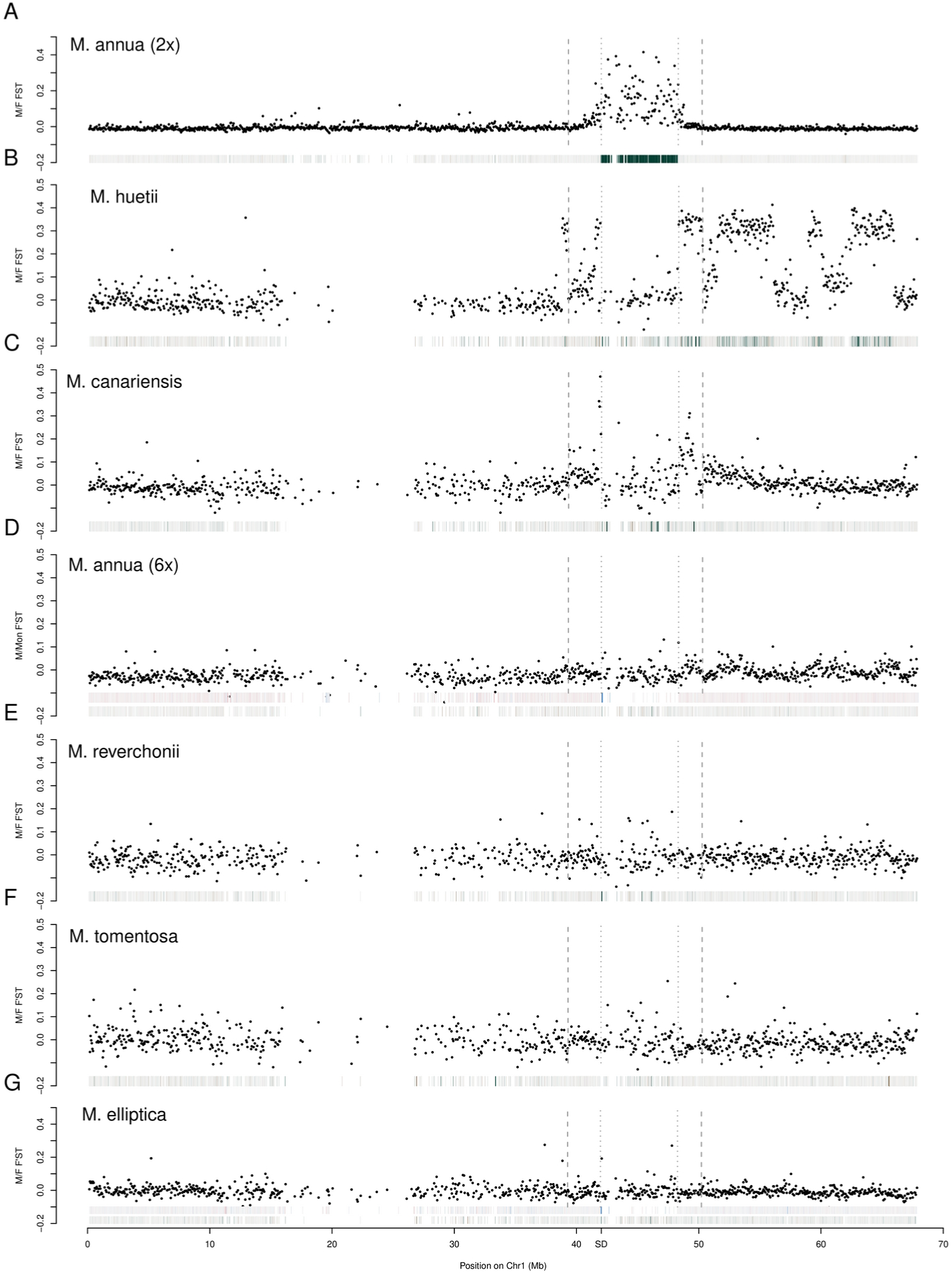
Differentiation and male-specific sequence content in different *Mercurialis* species. Manhattan plots show male-female *F*_ST_ in 50 kb windows in diploid *M. annua* (A) and *M. huetii* (B). In the allopolyploid lineagesspecies (C to -G) *F’*_ST_ is plotted instead. Bars at the bottom of the plot indicate male-specificity, with green probes being more male-specific. For hexaploid *M. annua* (D) and *M. elliptica* (G), an additional row of bars on top indicates male-specificity of gene-models based on PoolSeq data, with blue bars being more male-specific. Vertical dashed lines indicate the borders of the non-recombining region, dotted vertical lines indicate the borders of the old stratum.

Patterns from male-specific exon-capture probes show a strong enrichment of male-specific probes in the old stratum of diploid *M. annua* (Fig. 3A). Similar to male-female *F*_ST_ in *M. huetii*, regions with male-specific exon-capture probes extend beyond the SDR (Fig. 3B). In *M. canariensis*, fewer male-specific exon-capture probes are found in both the new and the old strata (Fig. 3C). In hexaploid *M. annua* and the three perennial species, fewer male-specific exon-capture probes are found in the old stratum of the SDR (Fig. 3D-G). Notably, male-specific exon-capture probes are found in all *Mercurialis* species at a position at the left edge of the old stratum, with the exception for *M. tomentosa*, where no exon-capture probes were found, likely due to insufficient sequence coverage. This region with shared male-specific exon-capture probes also overlaps with the gene model with the strongest male-specific sequencing coverage on Chromosome 1, based on PoolSeq data in hexaploid *M. annua* and *M. elliptica* (Fig 3D, Fig 3G).

## Discussion

### Two evolutionary strata on the sex chromosomes of *M. annua*

Our results point to two discrete nested regions on the sex chromosome of diploid *M. annua*, in which recombination between X and Y chromosomes has been suppressed at different times. The entire region is defined by a large inversion in the YY assembly (Fig. S1, Fig.1 D, Fig. 2A), which is found on the genome assembly and which is concordant with the lack of recombination on both male linkage maps (Fig. 1C Fig. 1E). In the centre of this larger region of reduced recombination is a second region that differs substantially from the flanking regions on both sides in terms of genetic degeneration and the accumulation of repeats (Fig 1A). Taken together these results suggest that recombination suppression in diploid *M. annua* evolved in two steps. The central region, which we call old stratum, evolved first, and a large inversion later extended the non-recombining region on both sides, formed a new, younger stratum. The lack of genetic degeneration, the low degree of synonymous divergence, and the limited accumulation of transposable elements in this younger stratum suggest that the inversion forming it must have been relatively recent, especially when compared to the old stratum. However, the greater number of premature stop codons identified based on reads from the YY assembly (Suppl. Table 1) suggests that the Y copy of the new stratum experiences reduced purifying selection.

While it is clear from our data that a single inversion formed the new stratum, it is unclear how recombination suppression evolved in the old stratum. Of the three gametologs that are still shared between X and Y sequences, two appear to have maintained the same order on the X and Y chromosomes while the remaining one did not. It is thus not clear whether structural variation was involved in initial recombination suppression. At the same time, there was a large accumulation of transposable elements in the old stratum of the Y, which increased its size by approximately 6 Mb. There also appear to have been multiple insertions of duplicated genes from the surrounding region, but also in two cases from other autosomes (Fig. 2A). There has also been substantial gene loss in the old stratum, with a total so that 22 genes found on the X chromosome but missing from the Y.

### Gene content in the old stratum

Among the genes missing from the Y chromosome of *M. annua*, one gene was annotated as DEFECTIVE TAPETUM AND MEIOCYTES 1 (DTM1). This gene is of particular interest, because it has been shown to be essential for tapetum development and meiosis in rice, and a knockout of the gene caused abnormal tapetal formation and developmental arrest of the development of pollen mother cells, causing male sterility (Ji et al. 2012). The similarity of the phenotypes of aborted pollen observed in rice with those of YY males in *M. annua* (Li et al. 2019) suggests that it is the lack of DTM1 on the Y chromosome that causes male sterility in YY males. This means that a gene that is essential for male fertility has been lost from the Y chromosome but is preserved on the X. Interestingly, the presence of such a recessive gene required for male function has been hypothesized to be involved in the gynodioecy model for the evolution of separate sexes in plants (Charlesworth & Charlesworth 1978). However, in that model the gene would be expected to be located on the (proto) Y chromosome and not the X chromosome, whereas here male sterility is only expressed in YY males, which are likely exceedingly rare in natural populations.

### Gene content in the new stratum

The new stratum consists of 387 genes, all of which are present in all three genome assemblies. With this relatively large number of genes, it is not surprising that some genes related to reproduction and sex expression should be included by chance. Indeed, we found several genes with functions related to male or female fertility and reproduction, including a tandem duplicate (SDR_NEW_G107, SDR_NEW_G108) of a gene, where the majority of sequences in the orthogroup were annotated as two-component response regulator ORR9. However, one of the sequences of the same orthogroup was annotated as ARR17, which acts as a master sex determination gene in Salicaceae (Leite Montalvão et al. 2022). (Suppl. Table 1). Among these genes there is a particularly interesting example potentially related to sex chromosome evolution. Five successive tandem copies (SDR_NEW_G15 – SDR_NEW_G19) were annotated as Reproductive Meristem 16 (REM16), and they are followed by an additional gene (SDR_NEW_G20) that we annotated as also belonging to the B3 domain containing family of REM genes. REM16 is involved in the regulation of flowering time in *Arabidopsis thaliana* (Yu et al. 2020) and in soy bean (Wang et al. 2020). In addition, REM16 showed strongly female-biased sex expression in dioecious *Cannabis sativa* and is considered a potential sex-determination gene (Shi et al. 2025). Like all other genes in the new stratum, the REM16 copies are found on all three genome assemblies, though we identified a premature stop codon in one of the copies on the YY genome assembly. If expression of REM16 confers an additive advantage to female reproductive fitness, it is conceivable that sexual selection could have favoured its increase in copy number. However, if such a fitness advantage caused a trade-off with male fitness, sexual conflict would arise, which in turn could have been mitigated by linkage of a non-functional allele to the male determining region via the observed inversion. Such a scenario would be consistent with the sexual antagonism model of recombination suppression (Rice 1987), and further analyses of gene expression and phenotypic effects of these genes in males and females could be used to test this hypothesis.

### Recombination suppression and male specific sequences in other *Mercurialis* lineages

Our comparisons of population genomic data *Mercurialis* points to variation in the size of the nonrecombining region and its degree of sequence divergence among species of the genus *Mercurialis*. Diploid *M. huetii* shows a strong signature of genetic differentiation between males and females, as well as the presence of male-specific exon-capture probes that overlap the non-recombining region that we defined here. However, this region extends beyond the non-recombining region by about 15 Mb to the right and, to a much smaller degree, also to the left. This suggests that there has been a substantial expansion of the non-recombining region in *M. huetii*, potentially as a result of additional structural variation.

In tetraploid *M. canariensis*, regions of increased differentiation (here measured as *F*‘_ST_) are found in both the old and new strata. Previous results suggested that *M. canariensis* inherited its Y chromosome from a close relative of diploid *M. annua* via allopolyploidization (Toups et al. 2022, Gerchen et al. 2022), consistent with our inference here of similar patterns of recombination suppression between the two species. Interestingly, our results point to a higher degree of differentiation in the new stratum of *M. canariensis* than in diploid *M. annua*. If our inferences are correct, then the new stratum in *M. annua and M. canariensis* should date to the same event, in which case the difference in X-Y differentiation between the two species might be related to demographic differences, e.g., a smaller effective population size of *M. canariensis*, which is currently only found in the Canary islands.

Our analysis of hexaploid *M. annua* and the three perennial species *M. reverchonii*, *M. tomentosa* and *M. elliptica* reveals windows with only moderately increased of *F*‘_ST_ in the sex-linked region or elsewhere in the genome. We cannot make clear inferences on the extent of the nonrecombining region in these species, but they suggest that at least no substantial expansion of the SDR occurred in them.

### APRR7 as putative sex master sex determination gene

Our analyses found a single gene, Arabidopsis Pseudo Response Regulator 7 (APRR7), that is male-specific in all *Mercurialis* species we compared. When comparing the location of male-specific probes in exon-capture datasets, we find a single male-specific exon-capture probe overlapping an exon of APRR7. In addition, our previously designed PCR probes, based on an early version of the *M. annua* genome assembly, also overlaps with APRR7 and shows male-specific amplification patterns in all *Mercurialis* species studied (Gerchen et al. 2022). Finally, our PoolSeq datasets point to APRR7 as having a male-specific pattern of sequence density in hexaploid *M. annua* and *M. elliptica* (Fig 3D, Fig. 3G). While also other genes show male-specific patterns of inheritance, depending on species and analysis, APRR7 is the only gene with consistent pattern of male-specific sequence variation across all *Mercurialis* species in our study (with the exception of *M. tomentosa*, where there is probably insufficient sequence coverage). We thus view it as a strong candidate for being a master sex-determination gene in dioecious and androdioecious species of the genus. APRR7 is part of the circadian clock pathway in *A. thaliana*, where it is also involved in the regulation of flowering time (Nakamichi et al. 2007). Orthologs of APRR7 have identified as targets of local adaptation in multiple plant species (Mayol et al. 2020, Whiting et al. 2024).

Our model of the evolution of sex chromosomes in *M. annua* would hint at an ancestral duplication of an autosomal copy of APRR7 on Chromosome 2. Neofunctionalization of this paralogue could have either directly caused the evolution of dioecy or a turnover of an existing sex chromosome system by acting as a dominant sex determination gene, which kept its functionality during multiple rounds of allopolyploidization in multiple annual and perennial *Mercurialis* species. Our results do not show any evidence for a second gene being involved in sex determination, raising the possibility that *Mercurialis* provides yet another example of single-gene sex determination in plants. Given that both males and females in *Mercurialis* occasionally produce fertile inflorescences of the opposite sex, implying that sterility mutations are not involved in sex determination, it seems plausible and indeed likely that separate sexes in the genus did not evolve via gynodioecy, as supposed by the canonical model for the evolution of dioecy (Charlesworth & Charlesworth 1978).

---

**Figure S1:**
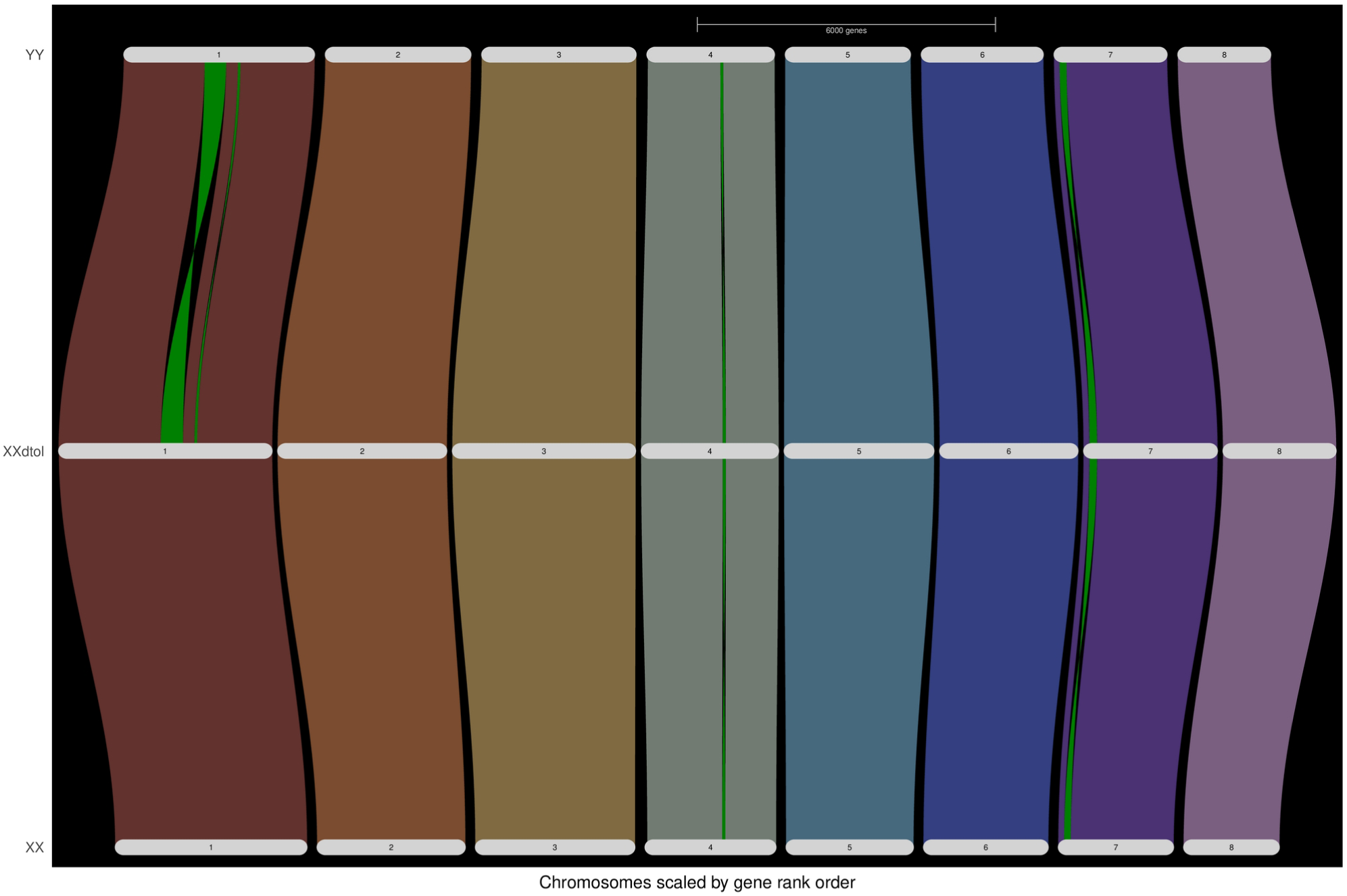
Riparian plot of synteny between genome assemblies inferred using GENESPACE. The eight colored bands represent chromosomes, the highlighted green regions represent inversions.

**Table S1:** Protein models used for Orthofinder3.

| **Species** | **RefSeq Genome version** |
| --- | --- |
| *Arabidopsis thaliana* | GCF_000001735.4_TAIR10.1 |
| *Cannabis sativa* | GCF_029168945.1_ASM2916894v1 |
| *Cucumis sativus* | GCF_000004075.3_Cucumber_9930_V3 |
| *Euphorbia lathyris* | GCF_963576675.1_ddEupLath1.1 |
| *Hevea brasiliensis* | GCF_030052815.1_ASM3005281v1 |
| *Manihot esculenta* | GCF_001659605.2_M.esculenta_v8 |
| *Populus trichocarpa* | GCF_000002775.5_P.trichocarpa_v4.1 |
| *Prunus persica* | GCF_000346465.2_Prunus_persica_NCBIv2 |
| *Quercus robur* | GCF_932294415.1_dhQueRobu3.1 |
| *Rosa chinensis* | GCF_002994745.2_RchiOBHm-V2 |
| *Ricinus communis* | GCF_019578655.1_ASM1957865v1 |
| *Vitis vinifera* | GCF_030704535.1_ASM3070453v1 |

